# The late endosome-resident lipid bis(monoacylglycero)phosphate is a cofactor for Lassa virus fusion

**DOI:** 10.1101/2021.03.22.436413

**Authors:** Ruben M. Markosyan, Mariana Marin, You Zhang, Fredric S. Cohen, Gregory B. Melikyan

## Abstract

Arenavirus entry into host cells occurs through a low pH-dependent fusion with late endosomes that is mediated by the viral glycoprotein complex (GPC). The mechanisms of GPC-mediated membrane fusion and of virus targeting to late endosomes are not well understood. To gain insights into arenavirus fusion, we examined cell-cell fusion induced by the Old World Lassa virus (LASV) GPC complex. LASV GPC-mediated cell fusion is more efficient and occurs at higher pH in cells expressing human LAMP1 compared to cells lacking this cognate receptor, but this receptor is not absolutely required for virus entry. GPC-induced fusion progresses through the same lipid intermediates as fusion mediated by other viral glycoproteins – a lipid curvature-sensitive intermediate upstream of hemifusion and a hemifusion intermediate downstream of acid-dependent steps that can be arrested in the cold. Importantly, GPC-mediated fusion is specifically augmented by an anionic lipid, bis(monoacylglycero)phosphate (BMP), which is highly enriched in late endosomes. We show that BMP promotes late steps of LASV fusion downstream of hemifusion – the formation and enlargement of fusion pores. This lipid also specifically promotes cell fusion mediated by GPC of the unrelated New World Junin arenavirus. The BMP-dependence of post-hemifusion stages of arenavirus fusion suggests that these viruses evolved to use this lipid as a cofactor to direct virus entry to late endosomes.

**Author Summary:** Pathogenic arenaviruses pose a serious health threat. The viral envelope glycoprotein GPC mediates attachment to host cells and drives virus entry *via* endocytosis and low pH-dependent fusion within late endosomes. Understanding the host factors and processes that are essential for arenavirus fusion may identify novel therapeutic targets. To delineate the mechanism of arenavirus entry, we examined cell-cell fusion induced by the Old World Lassa virus GPC proteins at low pH. Lassa virus fusion was augmented by the LAMP1 receptor and progressed through lipid curvature-sensitive intermediates, such as hemifusion (merger of contacting leaflets of viral and cell membrane without the formation of a fusion pore). We found that most GPC-mediated fusion events were off-path hemifusion structures and that the transition from hemifusion to full fusion and fusion pore enlargement were specifically promoted by an anionic lipid, bis(monoacylglycero)phosphate, which is highly enriched in late endosomes. This lipid also specifically promotes fusion of unrelated New World Junin arenavirus. Our results imply that arenaviruses evolved to use bis(monoacylglycero)phosphate to enter cells from late endosomes.

## Introduction

Old World (OW) and New World (NW) arenaviruses cause a range of diseases in humans, including severe hemorrhagic fever with high fatality rates of 15-35%. There are currently no FDA-approved vaccines or drugs to battle arenavirus infection. OW and NW arenaviruses infect a wide range of cells types *in vitro*, owing to their use of the ubiquitously expressed α-dystroglycan (α-DG) and transferrin receptor 1 (TfR1), respectively, for cell attachment and entry (reviewed in (1–3)). Binding of pathogenic Clade B NW arenaviruses to TfR1 initiates entry *via* clathrin-mediated endocytosis (3–5), whereas α-DG-driven OW arenavirus uptake occurs through a poorly characterized macropinocytosis-like pathway that is independent of clathrin, caveolin, dynamin-2, Rab5 and Rab7 (3, 5–9). Regardless of the specific receptor usage, NW and OW arenaviruses are thought to enter cells by undergoing low pH-triggered fusion with multivesicular bodies or late endosomes (2, 6, 8, 10, 11).

Fusion of arenavirus with host cells is mediated by the GP glycoprotein complex (GPC), a class I viral fusion protein (2, 3, 12–17). Like many viral glycoproteins, arenavirus GPC is synthesized as an inactive GPC precursor that is cleaved at two sites, one by a signal peptidase and the other by SKI-1/S1P protease. This generates a stable signal peptide (SSP) and non-covalently associated GP1 (surface) and GP2 (transmembrane) subunits. A unique feature of the arenavirus GP complex is that the SSP remains associated with the GP2 subunit after GPC cleavage. The ∼58-residue long SSP plays critical roles in GPC cleavage and transport to the plasma membrane and controls the initiation of GP conformational changes upon exposure to low pH (10, 18–23).

Interestingly, arenaviruses tend to rely on more than one host factor for entry. The Lassa Fever virus (LASV) switches from α-DG to the LAMP1 receptor in acidic endosomes (24–26). Similarly, a distant OW Lujo virus (LUJV) engages NRP-2 and CD63 (27). Efficient entry of the NW Junin virus has been reported to require TfR1 and a voltage-gated calcium channel (28). However, some arenaviruses can infect cells lacking the known receptors, albeit less efficiently (29–32). Although LAMP1 promotes LASV entry/fusion, it is not strictly required for the GPC fusion activity (31–33). Several lines of evidence indicate that acidic pH alone is sufficient to trigger LASV GPC conformational changes/functional inactivation (33), including GP1 dissociation from the GP2 (26) and fusion (31, 33) (but see (34) reporting LASV GPC resistance to acid treatment).

The molecular mechanism for arenavirus GPC-induced membrane fusion is not well understood. Progress in delineating the mechanism of arenavirus entry/fusion in late endosomes has been impaired by lack of knowledge regarding the precise pH or composition of intracellular compartments harboring the virus. Here, we used a cell-cell fusion model to examine the pH-, receptor-and lipid-dependence of LASV GPC-mediated fusion and characterize key intermediates of this process. We find that GPC-mediated cell-cell fusion is augmented by human LAMP1 expression and that GPC-LAMP1 binding shifts the pH optimum for fusion to a less acidic pH. We show that, similar to membrane fusion mediated by other viral proteins (e.g., (35–42)), LASV GPC-induced fusion progresses through a hemifusion intermediate and that fusion can be efficiently arrested upstream of hemifusion by incorporation of positive curvature-imposing lipids. Importantly, we find that GPC-mediated membrane fusion is specifically enhanced by a late endosome-resident lipid, bis(monoacylglycero)phosphate (BMP). These findings reveal BMP as a cofactor that drives efficient LASV fusion with late endosomes.

## Results

### GPC mediated membrane fusion is triggered by low pH and enhanced by LAMP1

We first examined LASV entry into cells using a luciferase reporter based single-round pseudovirus infection assay (see Methods). Human HEK293T and A549 cells were infected with HIV-1 luciferase reporter pseudoviruses bearing LASV GPC (referred to as LASVpp). Infection of both cell lines was pH-dependent, as evidenced by the abrogation of the luciferase signal in the presence of ammonium chloride that raises endosomal pH (Fig. 1A, third and sixth columns). LAMP1 knock down in these two cell lines using shRNA resulted in ∼2-3-fold reduction of LAMP1 expression (Fig. 1B) and reduced infection (Fig. 1A). The effect on infection was more pronounced in A549 cells expressing lower endogenous levels of LAMP1. In control experiments, LAMP1 knock down was without effect on infection by particles pseudotyped with the Influenza A virus hemagglutinin (IAVpp) in HEK293T cells, but enhanced IAVpp infection of A549 cells (Fig. 1A). The enhancing effect of LAMP1 knock down on IAVpp infection may be related to altered virus transport pathways and/or to changes in acidity of endosomal compartments. In agreement with the reported ability of LASV to enter cells lacking human LAMP1 in a pH-dependent manner (31, 32), LASVpp infected avian QT6 cells which express a LAMP1 ortholog that is not recognized by LASV GPC (25) (Fig. 1C). However, ectopic expression of human LAMP1 in these cells dramatically increased infection without significantly affecting IAVpp infection. The above results show that LAMP1 facilitates LASV entry into cells but is not absolutely required for GPC-mediated virus fusion with acidic endosomes.

**Figure 1.**
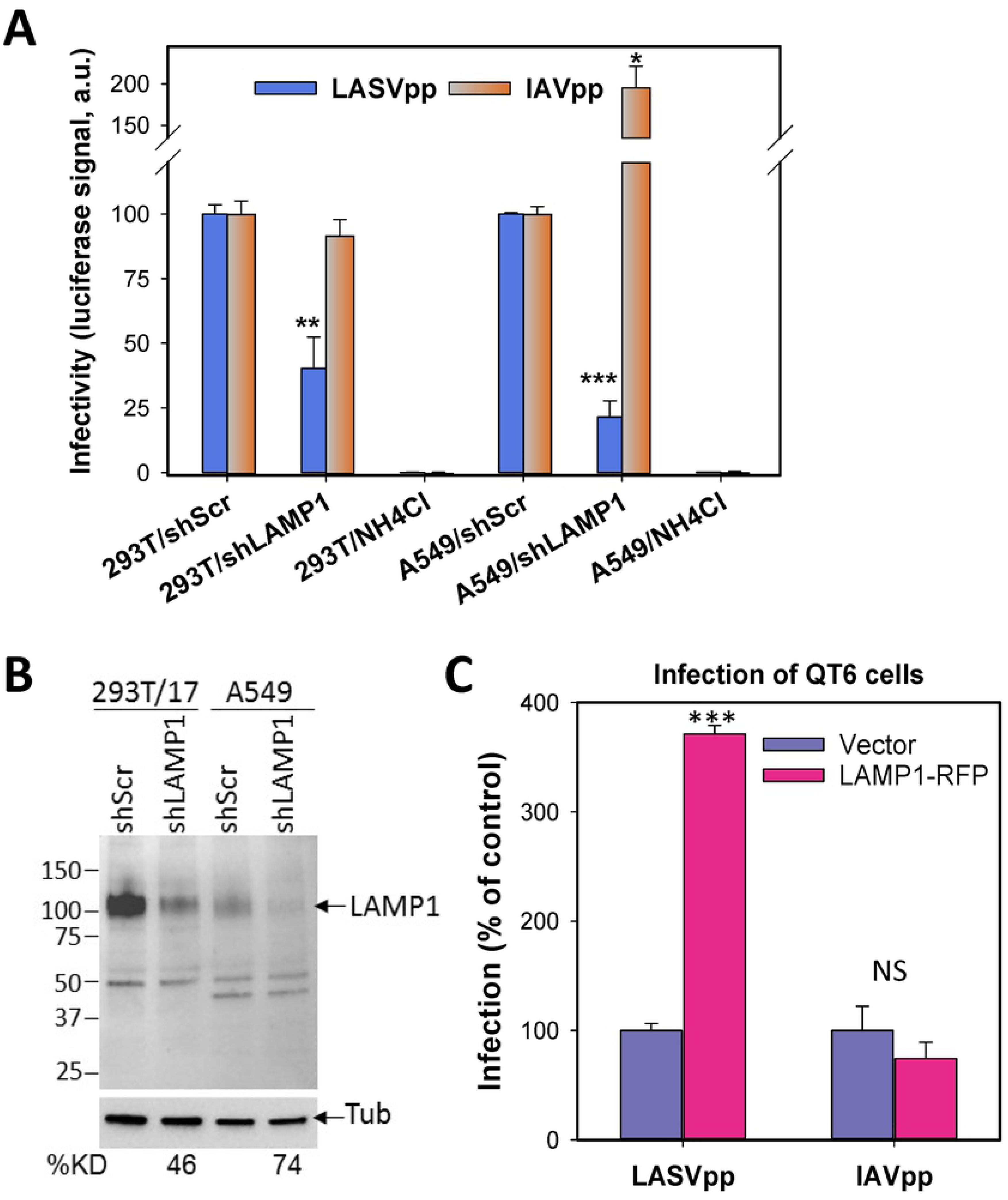
pH- and LAMP1-dependence of LASV pseudovirus infection. Human HEK2393T or A549 cells transduced with control (shScr) or LAMP1-specific shRNA (shLAMP1) were infected with luciferase-encoding HIV-1 particles pseudotyped with LASV GPC (LASVpp) or Influenza A HA/NA glycoproteins (IAVpp). Control infections were done in the presence of 30 mM NH_4_Cl to block virus entry from endosomes. The resulting luciferase signal was measured at 48 hpi and plotted on a log scale as mean and SEM from three independent experiments performed in duplicate. (B) Western blot analysis of LAMP1 expression and the efficiency of shRNA knockdown in HEK293T and A549 cells. Densitometry analysis of LAMP1 expression levels relative to control cells is shown under the bottom panel. Anti-tubulin served as the loading control. (C) LASVpp infection of avian QT6 cells transfected with an empty vector or human LAMP1-mRFP encoding plasmid. Shown are the results of a single experiment performed in triplicate. *, p < 0.05, **, p < 0.01, ***, p < 0.001, NS, not significant.

To further delineate the pH- and receptor-dependence of LASV GPC-mediated fusion, we employed a cell-cell fusion model that has been widely used for mechanistic studies of viral glycoprotein-mediated fusion (e.g., (31, 39, 43–48)). This model affords full control over extracellular pH, as well as access of fusion inhibitors to viral glycoproteins and cellular receptors throughout the fusion process. We transiently expressed LASV GPC in effector COS7 cells. Effector cells were brought in contact with target cells (HEK293T or avian QT6 or DF1 cells) transfected or not transfected with human LAMP1, and fusion was triggered by exposure to an acidic buffer. The resulting cell-cell fusion was assessed by fluorescence microscopy based on mixing of cytoplasmic dyes loaded into effector and target cells (e.g., Fig. 2A), as described previously (46).

**Figure 2.**
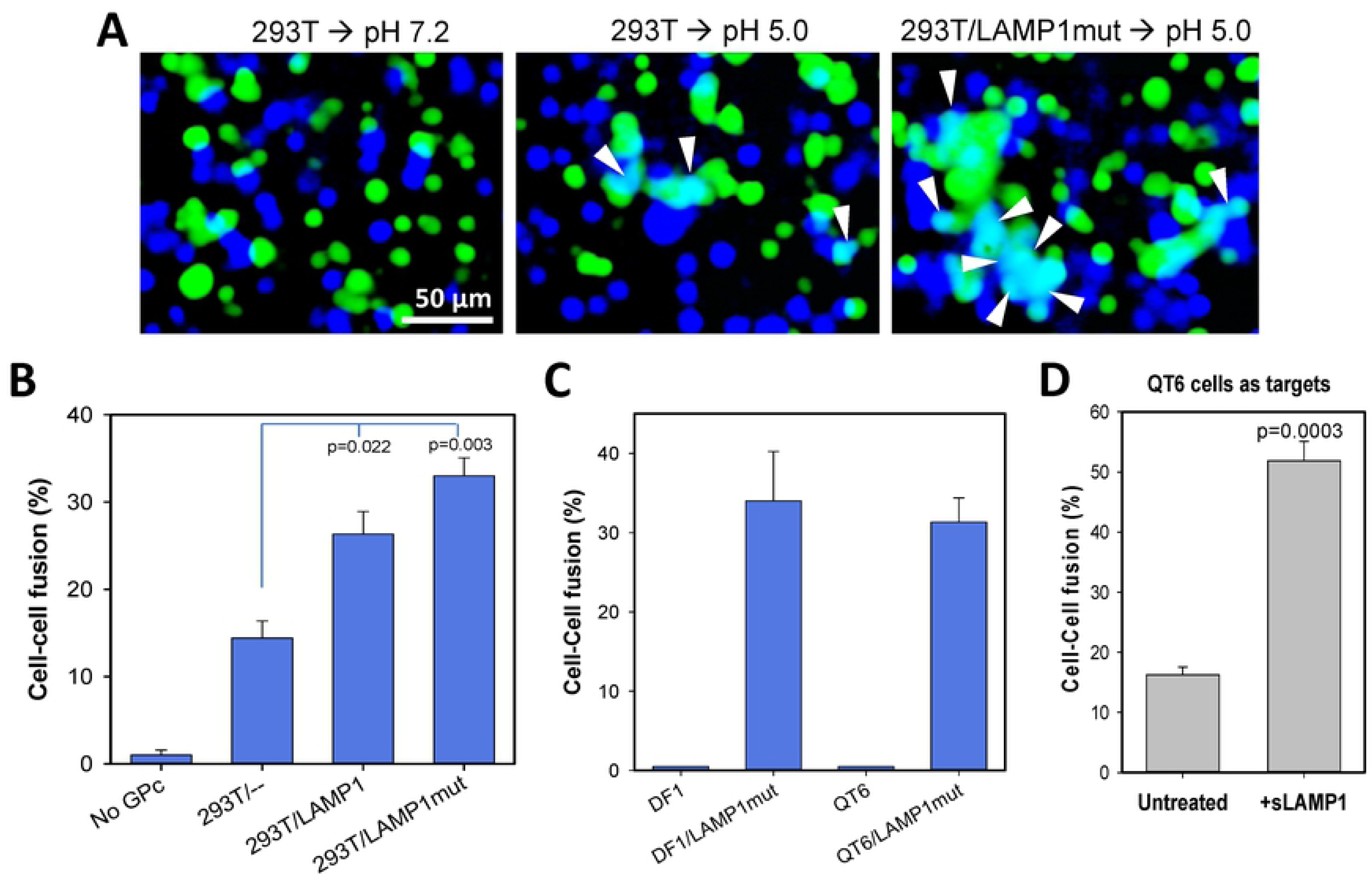
LAMP1-dependence of LASV GPC-mediated cell-cell fusion. (A) Representative images of effector COS7 cells transfected with LASV GPC and loaded with calcein-AM (green) and target HEK293T cells loaded with CMAC cytoplasmic dye (blue). HEK293T cells were transfected with the LAMP1 D384 mutant (LAMP1mut, right panel) or mock-transfected. Effector and target cells were mixed, allowed to adhere to poly-lysine-coated chambered slides and incubated for 30 min at room temperature. Fusion was triggered by exposing cells to a pH 5.0 buffer for 20 min at 37°C. Control wells were exposed to a pH 7.2 buffer (left panel). Double-positive fusion products are indicated by arrowheads. Scale bar 50 μm. (B) LAMP1-dependence of fusion between LASV GPC-expressing COS7 cells (loaded with calcein-AM) and HEK293T cells (loaded with CMAC) transfected with wild-type human LAMP1, LAMP1mut or mock-transfected. As a negative control, fusion between untransfected COS7 cells with HEK293T cells was measured. A 1:1 mixture of effector and target cells was adhered to poly-lysine coated coverslips and incubated 30 min at room temperature. Cell fusion was triggered by exposure to pH 6.2 for 10 min at room temperature (suboptimal trigger), and the fraction of cells positive for both cytoplasmic markers was measured after an additional incubation for 1 h at 37°C, neutral pH. The results are means and SEM from three independent experiments. (C) LAMP1-dependence of GPC-mediated fusion with avian DF-1 and QT6 cells. Avian cells transfected or not transfected with the human LAMP1mut were brought in contact with LASV GPC-expressing COS7 cells and exposed to pH 5.0 for 10 min at room temperature. The results are means and SEM from three independent experiments. (D) Soluble LAMP1 (sLAMP1) enhances GPC-mediated fusion with QT6 cells. GPC-expressing COS7 cells were co-plated with QT6 cells on coverslips for 30 min at room temperature to allow attachment and cell-cell contacts. The cells were further incubated in the absence or in the presence of 0.05 μg/ml of sLAMP1 for 20 min at room temperature, and fusion was triggered under optimal conditions (pH 5.0 at 37°C, 20 min). The results are means and SEM from three independent experiments.

We initially employed suboptimal conditions – pH of 6.2 and room temperature – to trigger Lassa GPC-mediated cell fusion. This pH is not sufficiently acidic to effectively trigger conformational changes in GPC that is not bound to the LAMP1 receptor (33), and should thus allow better appreciation of the LAMP1-dependence of LASV fusion. The modest level of GPC-mediated fusion with HEK293T cells under these suboptimal conditions (Fig. 2A, B) is in accord with a fraction of the endosome/lysosome-resident LAMP1 receptor expressed on the cell surface (49). Overexpression of LAMP1 or the LAMP1 D384 mutant (hereafter referred to as LAMP1mut) enhanced LASV GPC-mediated cell fusion (Fig. 2A, B). (LAMP1 mut lacks an endocytic signal contributing to better expression at the plasma membrane (25)). The observed mixing of two cytoplasmic markers was specifically induced by LASV GPC, since mock-transfected COS7 cells failed to fuse with target cells (Fig. 2B), and the arenavirus-specific inhibitor ST-193 suppressed fusion in a dose-dependent manner (Suppl. Fig. 1A).

Previous studies (31, 32) and our infectivity data (Fig. 1C) suggest that LAMP1 is not absolutely required for LASV fusion. To confirm the ability of LASV to undergo receptor-independent fusion, we fused GPC-expressing cells with avian DF-1 or QT6 cells expressing a LAMP1 ortholog that is not recognized by LASV (25). Whereas fusion with DF-1 or QT6 cells was not detected when a sub-optimal trigger for fusion was employed (pH 5.0, room temperature, Fig. 2C), an optimal trigger (pH 5.0, 37°C) or ectopic expression of the human LAMP1mut permitted highly efficient fusion (Fig. 2C, D). Lassa GPC fusion with QT6 cells was also markedly enhanced by addition of a soluble recombinant human LAMP1 (sLAMP1) (Fig. 2D). The above results show that, while not absolutely required for LASV GPC-mediated fusion, human LAMP1 markedly increases fusion efficiency.

### GPC-LAMP1 interaction allows LASV fusion at higher pH

We assessed the pH-dependence of LASV fusion by exposing the effector/HEK293T cell complexes to buffers of different acidity. A pH threshold of LASV GPC-mediated cell fusion was ∼6.2, with maximal fusion at pH 4.8 (Fig. 3A), in general agreement with previous studies (10, 31). Ectopic expression of LAMP1mut in target cells increased the overall efficiency of fusion, while not significantly affecting the shape of the curve (Fig. 3A). The modest reduction in fusion efficiency at pH<4.8 may be caused by acid-mediated GPC inactivation, which may be caused by shedding of the GP1 subunit of GPC, known to occur at pH ≤ 4.0 in the absence of a cognate receptor (Suppl. Fig. S1B and (26, 33)). Parallel experiments with avian QT6 cells lacking human LAMP1 revealed a lower extent of fusion and an overall shallower pH-dependence compared to QT6 cells transfected with LAMP1mut (Fig. 3B). Also, the pH-dependence of fusion with QT6 cells did not reach a plateau at pH 4.8, regardless of LAMP1mut expression (Fig. 3B). The greatly increased fusion with LAMP1mut-expressing QT6 cells at mildly acidic pH supports the notion that LAMP1 binding shifts the pH optimum of GPC-mediated fusion to higher values (25, 31, 33). A linear pH-dependence of fusion with both QT6 and QT6/LAMP1mut cells at pH below ∼6.0 suggests that this fusion is predominantly acid-dependent, but receptor-independent. The difference between the pH-dependence of fusion with HEK/293T and QT6 cells is surprising and might indicate that cell type- or species-specific factors other than LAMP1 promote LASV GPC refolding at acidic pH in the absence of the receptor.

**Figure 3.**
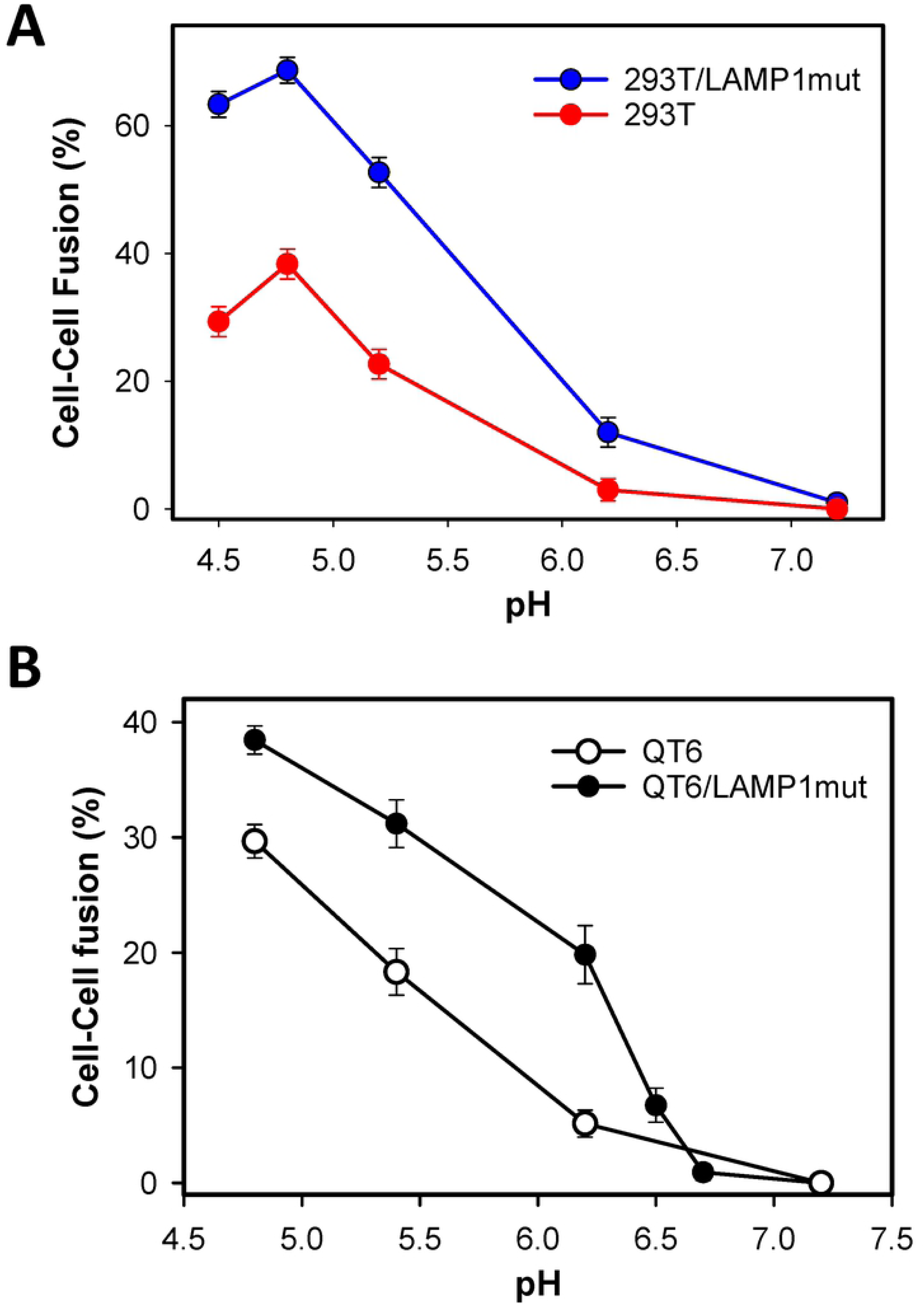
pH-dependence of LASV GPC-mediated cell-cell fusion. (A) Fusion between COS7 cells transfected with LASV GPC and HEK293T cells mock-transfected or transfected with LAMP1mut was triggered by exposure to buffers of different acidity for 20 min at 37°C (optimal trigger). The results are means and SEM from three independent experiments normalized to the maximal fusion values observed at pH 4.8. (B) pH-dependence of fusion, using the protocol described in panel A, was measured between LASV GPC-expressing COS7 cells and plain QT6 cells or QT6 cells transfected with LAMP1mut. The results are means and SEM from three independent experiments normalized to the value at the lowest pH.

### LASV fusion is reversibly arrested by positive-curvature lipids

Diverse membrane fusion reactions can be blocked by exogenous lyso-lipids that confer positive curvature to the contacting membrane leaflets and thereby prevent the formation of a net negative curvature stalk structure (Fig. 4A and (50–52)). This lipid-arrested stage (LAS) is largely reversible upon washing away lysolipids and commences at neutral pH (44, 53), implying that the viral proteins are arrested in a “committed” stage that is downstream of low pH-dependent steps. Like other protein-mediated fusion reactions, LASV GPC-mediated fusion was blocked when lyso-PC was present before and during a low pH pulse. Virtually no cell-cell fusion was observed while lyso-PC was present (Fig. 4B), but fusion ensued upon removal (at neutral pH) of this lipid (Fig. 4B). This result implies that LASV fusion progresses through curved lipid intermediate(s) of net negative curvature, likely a stalk and a hemifusion diaphragm.

**Figure 4.**
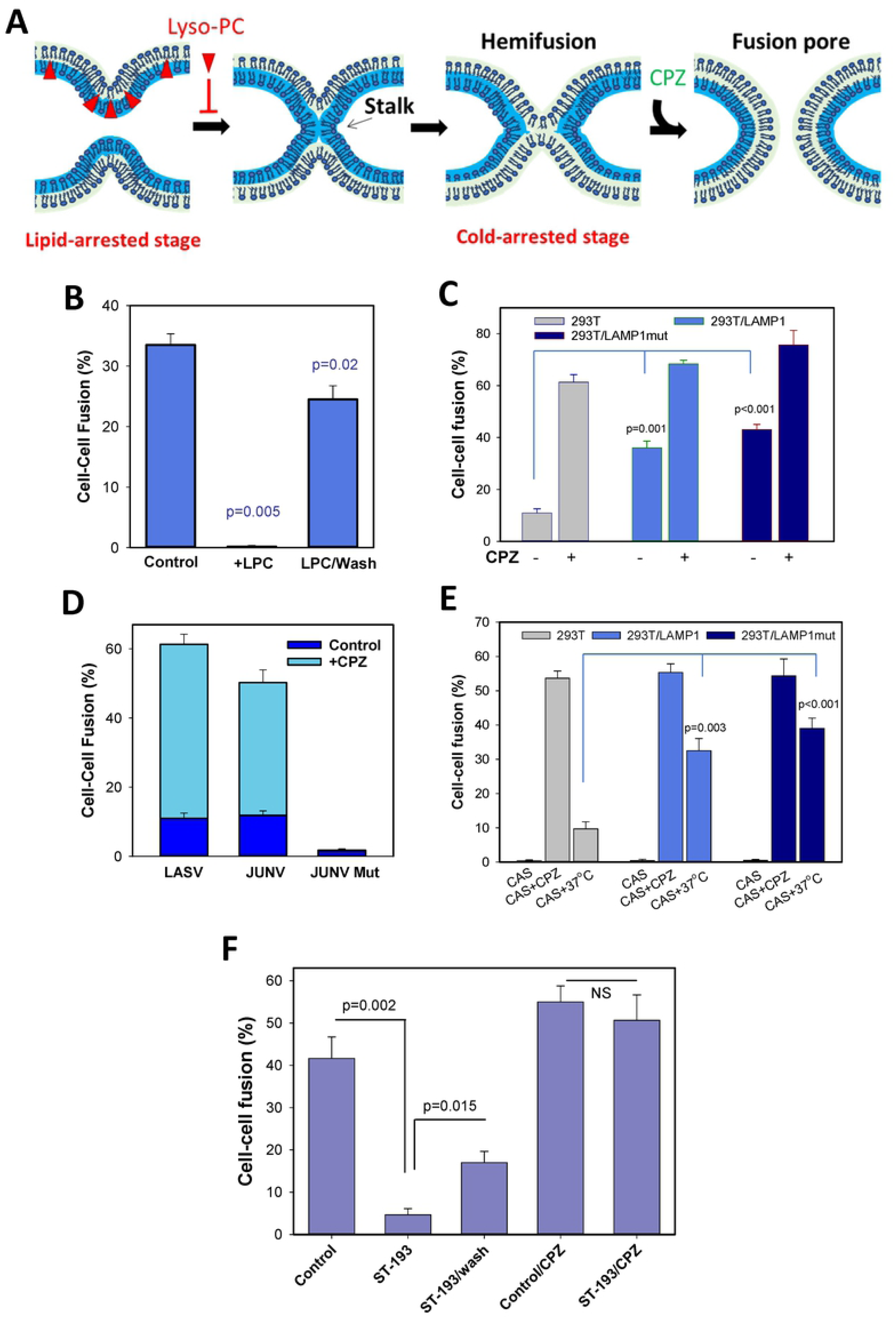
LASV GPC-mediated fusion progresses through a hemifusion intermediate that is blocked by lyso-lipids. (A) Illustration of cold-arrested (CAS, hemifusion) and lyso-lipid-arrested (LAS, pre-hemifusion) intermediates and the effect of chlorpromazine (CPZ). (B) Lipid-arrested stage of LASV fusion. GPC-expressing COS7 cells were bound to QT6 cells transfected with LAMP1mut in the presence of 285 μM stearoyl lyso-PC for 30 min at room temperature, followed by exposure to pH 5.0 at room temperature in the presence of lyso-PC. Cells were either washed with delipidated BSA (2 mg/ml) to remove lyso-PC (right bar) or left with lyso-PC (middle bar) and incubated for 30 min at 37°C. The results are means and SEM from three independent experiments. (C) The effect of CPZ on fusion between GPC-expressing COS7 cells and HEK293T cells mock-transfected or transfected with LAMP1 or LAMP1mut. Cell fusion was triggered under sub-optimal conditions (pH 6.2, 10 min at room temperature). After an additional 1 h incubation at 37°C, cells were treated with CPZ (0.5 mM) for 1 min or left untreated. The results are means and SEM from three independent experiments. (D) CPZ enhances LASV and Junin GPC-mediated cell fusion. Fusion between LASV or Junin GPC-expressing COS7 cells and HEK293T cells was triggered by exposure to pH 6.2 (for LASV) or pH 5.5 (JUNV). After a 20 min incubation at 37°C, cells were treated for 1 min with 0.5 mM CPZ at room temperature (n=3). (E) Cold-arrested intermediate (CAS) of LASV GPC fusion. GPC-expressing COS7 cells were incubated with HEK293T cells (transfected or not with LAMP1 or LAMP1mut) for 30 min at room temperature and exposed to pH 6.2 on ice for 10 min. Cells were either immediately treated with cold 0.5 mM CPZ for 1 min or additionally incubated for 1 h at 37°C, neutral pH. The results are means and SEM from three independent experiments. (F) ST-193 captures LASV fusion at a hemifusion intermediate. COS7 cells expressing LASV GPC were mixed with HEK293T cells, attached to glass slides and allowed to adhere and form contacts for 30 min at room temperature. Fusion was initiated by exposure to pH 5.0 for 15 min at 37°C, in the presence or absence of 150 μM ST-193. Cells were either washed to remove the inhibitor or kept in the presence of ST-193, and either immediately treated with 0.5 mM CPZ for 1 min or further incubated at neutral pH for 1 h, 37°C. The results are means and SEM from three independent experiments.

### LASV fusion progresses through a hemifusion intermediate that can be arrested at low temperature

Membrane fusion mediated by diverse viral glycoproteins progresses through a common hemifusion intermediate (reviewed in (54–56)) (Fig. 4A). This intermediate is formed through merger of the contacting leaflets of two membranes, while distal leaflets form a new bilayer referred to as hemifusion diaphragm (57, 58) (Fig. 4A). It has been shown that low pH-triggered viral fusion can be arrested at a local hemifusion stage in the cold (referred to as cold-arrested stage, CAS) (46, 53, 59–61) (Fig. 4A). After creating CAS, fusion can be recovered at neutral pH by raising temperature. In other words, steps of fusion downstream of CAS are temperature-dependent but no longer require low pH. The formation of a hemifusion intermediate, including the one formed at CAS, can be indirectly inferred by treating cells with chlorpromazine (CPZ) at neutral pH, which selectively destabilizes a hemifusion diaphragm and promotes full fusion (57) (Fig. 4A).

We first asked if LASV GPC-mediated fusion exhibits a natural tendency to get stuck at a hemifusion stage, as is the case for membrane fusion mediated by other low pH-dependent viruses, especially when suboptimal triggers for fusion (insufficiently low pH and/or reduced temperature) are employed (44, 53, 61). We found that GPC-mediated fusion triggered under suboptimal conditions (pH 6.2, room temperature) was markedly enhanced by a brief exposure to CPZ after returning to neutral pH (Fig. 4C). This CPZ-mediated fusion enhancement was much more pronounced for plain HEK293T cells (>6-fold), as compared to cells expressing wild-type or mutant LAMP1 (∼2-fold, Fig. 4C). Notably, over-expression of LAMP1 markedly decreased the fraction of dead-end hemifusion structures: structures that did not naturally progress to full productive fusion. Higher levels of LAMP1 on cell surfaces did not considerably increase the overall formation of hemifusion sites, but markedly enhanced the probability of transition to full fusion at 37°C. A similar dramatic increase in cell-cell fusion was observed after treatment of Junin virus GPC-mediated cell fusion products with CPZ, whereas this treatment was without effect in control experiments using cells expressing a fusion-incompetent GPC mutant (20) (Fig. 4D). Thus, under suboptimal conditions, GPC tends to create hemifusion structures that do not resolve into full fusion; these structures can be forced to convert to full fusion by CPZ treatment.

Next, we tested the possibility of capturing LASV fusion at a cold-arrested stage (CAS, Fig. 4A) by exposing effector/target cell pairs to low pH in the cold. A reversible arrest of GPC-mediated cell-cell fusion at CAS was evident by the commencement of fusion after shifting to 37°C at neutral pH (Fig. 4E). As observed previously for uninterrupted LASV GPC-mediated cell-cell fusion (Fig. 2), over-expression of LAMP1 or LAMP1mut markedly enhanced the extent of fusion after shifting cells captured at CAS to 37°C (Fig. 4E). Importantly, CPZ treatment of cells arrested at CAS (after returning to neutral pH, still at low temperature) promoted efficient fusion (Fig. 4E), consistent with the formation of local hemifusion structures at CAS. In control experiments, exposure to CPZ without low pH pretreatment did not result in significant cell-cell fusion (data not shown). CPZ treatment induced more fusion between cells captured at CAS than a shift to 37°C and this difference was more apparent for fusion with suboptimal target cells (Fig. 4E). These results show that GPC-LAMP1 interactions increase the probability of conversion of hemifusion to full fusion.

### Arenavirus fusion inhibitor captures LASV GPC-induced fusion at a hemifusion stage

The small-molecule arenavirus fusion inhibitor ST-193 has been shown to block pH-induced conformational changes in GPC, including the shedding of the GP1 subunit, but the fusion block can be overcome at sufficiently acidic pH (62). ST-193 is thought to act by binding to the interface between the GP2 subunit transmembrane domain and the stable signal peptide (62) and disfavoring the early steps of GPC refolding at low pH. Since, the compound appears to interfere with pH-induced GPC refolding, we asked whether ST-193 blocks of LASV fusion at early steps prior to membrane merger. Effector-target cell complexes were exposed to pH 5.0 at 37°C in the presence of a high concentration ST-193 that almost completely inhibits cell-cell fusion (Fig. 4F and Suppl. Fig. S1A). Subsequent removal of the inhibitor at neutral pH resulted in a partial recovery of fusion (Fig. 4F). This partial reversibility of LASV fusion indicates that a large fraction of GPC proteins undergoes irreversible conformational changes and inactivates at low pH in the presence of ST-193. Importantly, CPZ application immediately after removal of ST-193 and in control samples not exposed to ST-193 induced efficient fusion that exceeded the level of uninterrupted fusion at 37°C (Fig. 4F). These findings show that, in the presence of ST-193, GPC undergoes low pH-dependent conformational changes that promote membrane hemifusion. Similar levels of CPZ-mediated fusion for control and ST-193 treated samples (Fig. 4F) suggest that acid-dependent steps of GPC-mediated membrane fusion are not grossly affected by this inhibitor when cells are subjected to an optimal trigger. However, in the presence of ST-193, GPC primarily creates dead-end hemifusion structures that can only be converted to full fusion by CPZ treatment; less than 30% of the hemifusion structures naturally resolves into full fusion at 37°C.

### Late steps of LASV GPC-mediated fusion are enhanced by a late endosome-resident lipid

LASV is thought to fuse with multivesicular bodies/late endosomes (2, 3, 6, 8). However, LASV GPC can mediate fusion with LAMP1-expressing cells at mildly acidic pH typical for early endosomes (Fig. 3 and (31–33)). We therefore asked if LASV entry from early endosomes may be delayed due to the need for an additional host factor localized to late endosomes. The unusual anionic lipid, bis(monoacylglycero)phosphate (BMP), also known as lysobisphosphatidic acid, is greatly enriched in late endosomes (around 15% of the total lipids (63, 64)). In fact, preincubation of cells with anti-BMP antibodies has been reported to diminish LASV infection (6). However, prolonged incubation with anti-BMP antibodies is known to disrupt several essential cellular processes, including cholesterol transport and biogenesis of multivesicular bodies (65–68). These pleotropic effects may indirectly disfavor LASV fusion by disrupting virus uptake and/or transport to permissive intracellular compartments. In this regard, cell-cell fusion appears well-suited for exploring lipid-dependence of LASV GPC-mediated fusion.

To investigate the effect of lipids on LASV fusion, the effector/target cell complexes were treated with BMP or zwitterionic lipid DOPC (control) prior to exposure to low pH. We observed specific promotion of LASV GPC-mediated cell fusion by exogenous BMP, but not by DOPC (Fig. 5A). This enhancing effect of BMP was observed across the range of LAMP1 expression, using plain HEK293T cells and cells transfected with LAMP1 or LAMP1 mutant (Fig. 5A). Also, BMP, but not another anionic lipid, DOPS, markedly enhanced GPC-mediated fusion with suboptimal QT6 cells (Fig. 5B). BMP also enhanced the Junin virus GPC-mediated cell fusion by ∼4-fold, whereas DOPC and DOPS were without effect (Fig. 5C). In contrast, influenza HA-mediated fusion was not affected by either BMP or DOPC, but was modestly promoted by DOPS. Note that failure of exogenous DOPS to promote cell-cell fusion mediated by GPC was not due to the lack of incorporation into the plasma membrane, as revealed by staining cells with Annexin V (Suppl. Fig. S2A).

**Figure 5.**
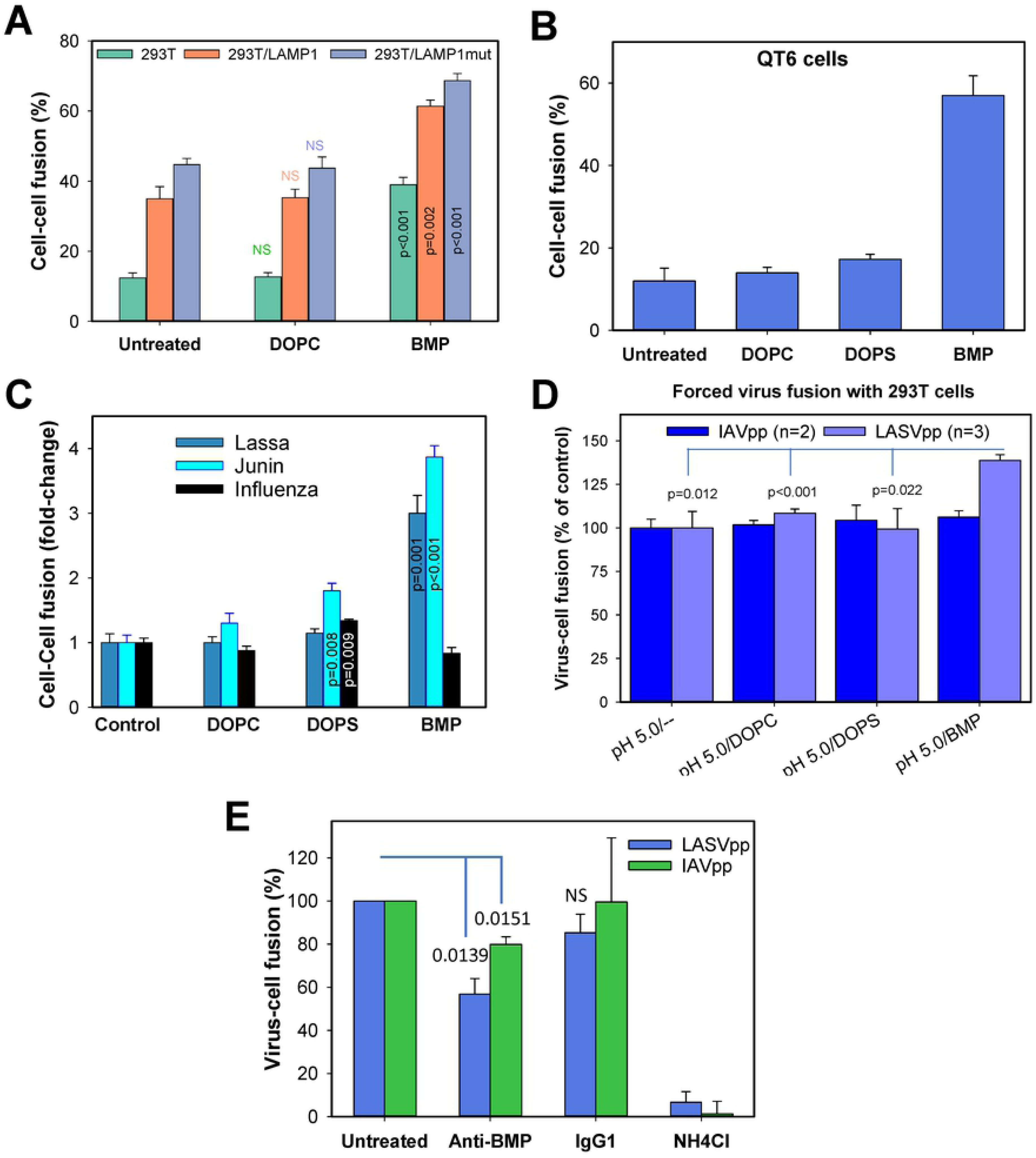
Lipid-dependence of LASV fusion. (A) COS7 cells transiently expressing LASV GPC were co-incubated with mock-transfected or LAMP1 or LAMP1mut transected 293T cells in the presence of 10 μg/ml DOPC or BMP dissolved in BSA (1 mg/ml) for 20 min at room temperature. Cell fusion was triggered by exposure to pH 6.2 at room temperature in the absence of exogenous lipids and was measured after an additional incubation for 1 h at 37°C. The results are means and SEM from three independent experiments. (B) GPC-expressing COS7 cells were incubated with QT6 cells in the presence of 10 μg/ml DOPC, DOPS or BMP dissolved in BSA (1 mg/ml) for 20 min at room temperature. Cell fusion was triggered by exposure to pH 5.0 at 37°C for 20 min in the absence of exogenous lipids. The results are means and SEM from three independent experiments. (C) COS7 cells expressing LASV or Junin Virus GPC or IAV HA were brought in contact with HEK293T cells, incubated with 10 μg/ml DOPC, DOPS or BMP dissolved in BSA for 20 min at room temperature and exposed to pH 5.0 for 20 min at 37°C. The results are means and SEM from four independent experiments. (D) Lipid-dependence of forced virus-cell fusion. LASVpp or IAVpp were added to HEK293T cells pretreated with 0.2 μM BafA1 and incubated for 1h at 37°C prior to treatment with 40 μg/ml of indicated lipids dissolved in BSA. Virus fusion to cells was induced by exposure to pH 5.0 for 30 min at 37°C in the presence of lipids. The cells were next loaded with the BlaM substrate and the extent of virus-cell fusion measured after overnight incubation, as described in Methods. The results are means and SEM from 3 independent experiments (LASVpp) and two independent experiments (IAVpp). (E) Human A549 cells were starved for 6 h and incubated with 50 μg/ml of anti-BMP or control IgG1κ antibodies for 15 h before infecting with LASVpp or IAVpp. Virus-cell fusion was measured using a BlaM assay. Control samples were treated with 70 mM NH_4_Cl to block endosomal entry of viruses. The results are means and SD from 2 independent experiments performed in duplicates.

In contrast to the effect of BMP on cell-cell fusion, a direct LASV pseudovirus-cell fusion assay based on a beta-lactamase reporter (69) did not reveal a significant effect of this or other lipids (Suppl. Fig. S3). A possible reason for the lack of BMP effect on viral fusion is that late endosomes already contain high levels of this lipid, so exogenous addition of BMP does not significantly alter their composition. To address this possibility, we first tested the lipid-dependence of LASVpp entry by forcing its fusion with the plasma membrane, which does not contain BMP (70). LASVpp fusion with the plasma membrane was triggered through exposure to low pH in the presence of Bafilomycin A1, which, by raising endosomal pH, blocks the conventional route of virus infection. This forced fusion protocol revealed that pretreatment of virus-cell complexes with BMP, but not DOPC or DOPS, significantly increases the extent of virus fusion with the plasma membrane (Fig. 5D). By contrast, forced virus-cell fusion mediated by the IAV HA glycoprotein was not affected by BMP. In parallel experiments, we assessed the ability of anti-BMP antibodies to interfere with LASVpp entry through an endocytic pathway, as reported previously (6). Overnight pre-treatment of A549 cells with anti-BMP, but not with control isotype antibodies, inhibited LASVpp fusion and, to a lesser degree, IAVpp fusion (Fig. 5E). The modest effect on IAVpp fusion may be related to disruption of multivesicular body biogenesis and/or of cholesterol transport (65–68). The above results imply that arenavirus GPC-mediated membrane fusion is specifically promoted by BMP and that this dependence may favor LASV entry from late endosomes.

### BMP promotes the formation and growth of GPC-mediated fusion pores

To further delineate the role of BMP in GPC-mediated fusion, we asked whether the effect is sensitive to whether this lipid is incorporated in a target, in GPC-expressing membranes, or both. Effector or target cells detached from culture dishes were separately treated with DOPC or BMP, mixed and exposed to low pH. In control experiments, a mixture of effector and target cells was pretreated with indicated lipids. As shown in Suppl. Fig. 2B, addition of BMP increased the efficiency of GPC-mediated cell fusion by ∼2.5-fold compared to DOPC control, irrespective of whether BMP was incorporated into effector or target cells. The ability of BMP to promote fusion upon incorporation into effector cells may indicate direct interaction with GPC or its effect on late, post-hemifusion steps of fusion whereupon the contacting leaflets of two membranes are merged.

We asked whether BMP augments early, low pH-dependent steps of GPC-mediated fusion or plays a role in downstream, pH-independent steps of viral fusion. Two strategies were used to address this question. First, the products of GPC-mediated fusion of cells pretreated with BMP or DOPC or left untreated (negative controls) were briefly exposed to CPZ to fully fuse cells arrested at hemifusion. The increase in fusion induced was dramatic for untreated or DOPC-pretreated cells, but modest for BMP-pretreated cells which had largely fused prior to addition of CPZ (Fig. 6A). The extent of fusion subsequent to CPZ addition was independent of pretreatment. These results suggest that the efficiency of formation of CPZ-sensitive hemifusion structures is independent of BMP, whereas the probability of conversion of these intermediates into full fusion is dramatically and specifically increased by BMP. The second strategy was to capture cell-cell fusion at CAS, add various lipids, at neutral pH, and then raise temperature. Fusion was potently promoted by the addition of BMP at CAS, but not by the addition of DOPC or DOPS (Fig. 6B).

**Figure 6.**
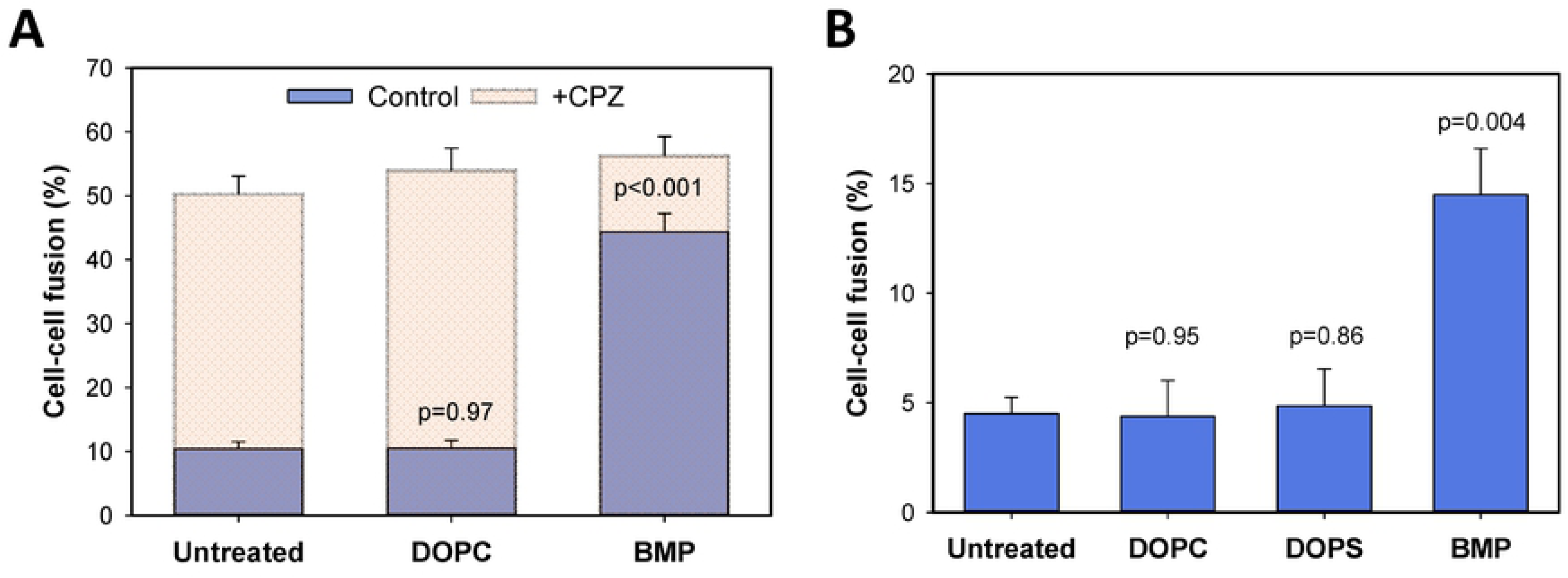
BMP promotes LASV fusion at a stage downstream of hemifusion. (A) BMP facilitates transition from hemifusion to fusion. COS7 cells expressing LASV GPC were incubated with HEK293T cells in the presence of 10 μg/ml DOPC or BMP dissolved in BSA (1 mg/ml) for 20 min at room temperature. Cell fusion was initiated by exposure to pH 6.2 for 10 min at room temperature in the absence of lipids followed by incubation at 37°C for 1 h. followed by a brief (1 min) exposure to 0.5 mM CPZ or to PBS (control). The results are means and SEM from three independent experiments. (B) Lipid-dependence of transition from cold-arrested stage (CAS) to full fusion. Fusion between LASV GPC-expressing COS7 cells and QT6 cells was initiated by exposure to pH 5.0 for 10 min at 4°C followed by treatment with 10 μg/ml of indicated lipids in BSA for 10 min at room temperature, at neutral pH, and an additional 30 min-incubation at 37°C.

Finally, we examined the effect of exogenously added BMP on the size of GPC-mediated fusion pores and their propensity to enlarge. Toward this goal, we performed time-lapse imaging of the redistribution of a small cytoplasmic marker, calcein, between effector and target cells (Fig. 7A, B) and deduced the pore permeability from the rate of dye redistribution, as described previously (71, 72). Interestingly, GPC-mediated redistribution of calcein between the dye donor and acceptor cell pairs was much slower than for cell-cell fusion mediated by other viral glycoproteins (43, 71)). Moreover, calcein redistribution often stalled after a few minutes so that the dye did not fully equilibrate between the two cells (Fig. 7A, B). This result suggests that GPC-mediated fusion pores remain small under our experimental conditions and even tend to close. The average pore permeability profile confirmed the failure of GPC-mediated fusion pores to grow (Fig. 7E). In stark contrast, fusion pores formed between BMP-pretreated cells grew efficiently, as evidenced by a quick and complete redistribution of calcein (Fig. 7C, D and E). Control experiments showed no significant effect of DOPC on the initial size or enlargement of GPC-mediated fusion pores (data not shown and Fig. 7E). We also measured the kinetics of fusion pore formation based on the onset of calcein redistribution and found that BMP pretreatment markedly accelerated the rate of cell-cell fusion (Fig. 7F). Collectively, the above results show that BMP selectively enhances post-hemifusion steps of LASV fusion, including the formation and dilation of fusion pores.

**Figure 7.**
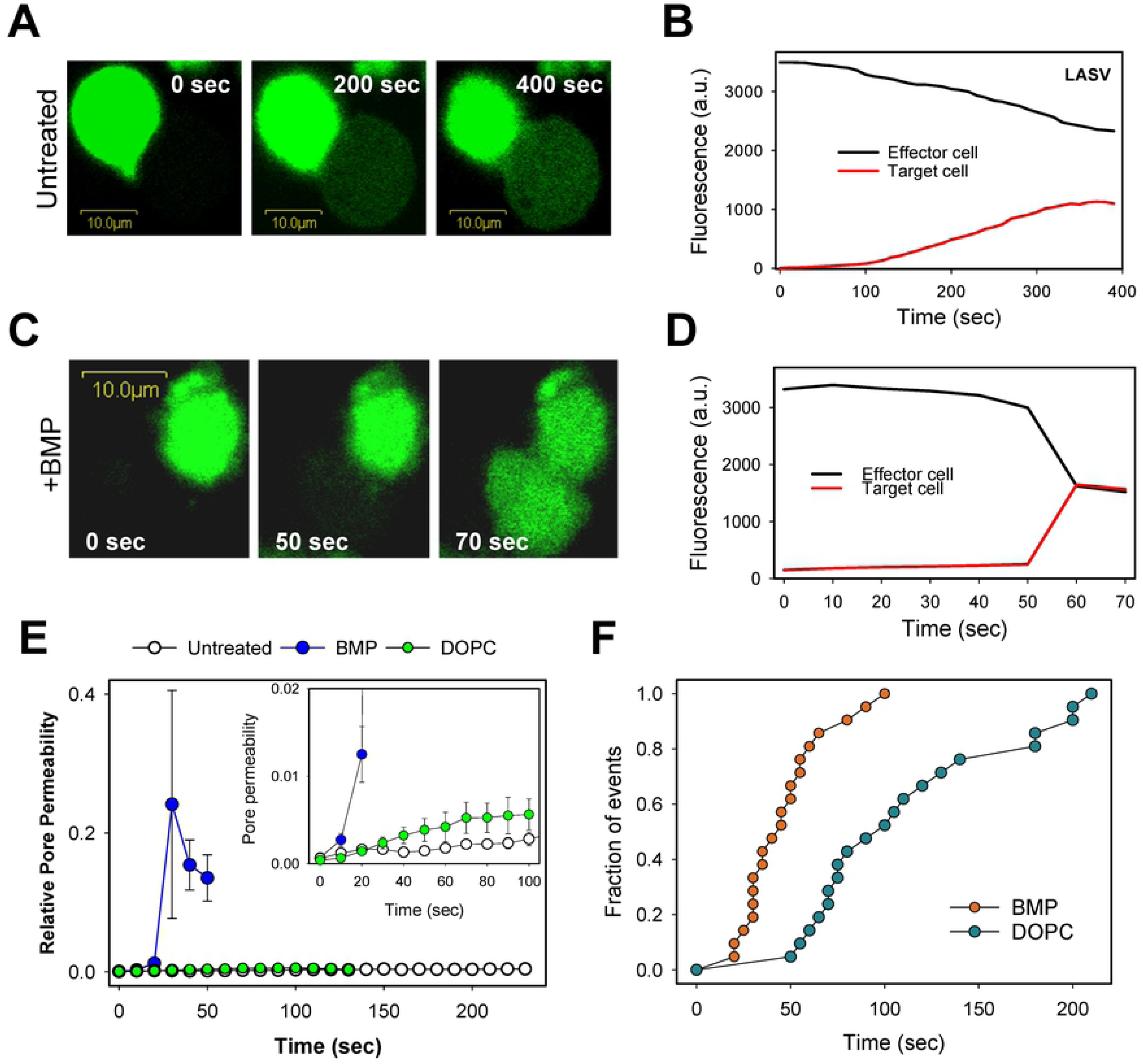
BMP markedly promotes LASV GPC-mediated fusion pore formation and enlargement. Effector cells loaded with calcein (green) were mixed with unlabeled target cells and adhered to poly-lysine coated coverslips. Cells were then pre-treated with 20 μg/ml of BMP or DOPC for 10 min at room temperature or left untreated. Cell-cell fusion was triggered by transferring cells to a pH 5.0 buffer and quickly raising the temperature to 37°C through an IR temperature jump protocol (see Methods). (A, B) Snapshots and fluorescence intensities showing partial calcein redistribution between untreated effector and target (dye donor and acceptor) cells. (C, D) Same as in panels A, B, but for cells pretreated with BMP. (E) Ensemble average of permeabilities of six fusion pores for untreated and DOPC- or BMP-treated cells. Traces from different pores were aligned so that t = 0 represents a time point immediately before dye movement was detected. *Inset*: Initial permeability profiles of fusion pores. Error bars are SEM. (F) Early kinetics of fusion pore formation for control and BMP-treated cells. The data points represent time after raising the temperature until the onset of calcein transfer.

## Discussion

Here, using single-cycle infectivity and cell-cell fusion assays, we show that, in agreement with published work (25, 31, 32), LASV GPC-mediated fusion is triggered by low pH and augmented by interaction with human LAMP1. Further, we find that, like fusion mediated by other viral proteins, LASV GPC-induced fusion progresses through highly curved lipid intermediates – stalk and a downstream hemifusion structure that are strongly modulated by the lipid composition of the fusing membranes. However, in spite of common features shared with other viral fusion proteins, arenavirus GPC-mediated fusion exhibits several unique properties: (1) resistance to restriction by interferon-induced transmembrane proteins (IFITMs) (35, 73, 74); (2) GPC-mediated permeabilization of the viral membrane prior to fusion (73); and (3) dependence on BMP for efficient fusion (Figs. 5-7).

We also show that LASV GPC-mediated fusion with avian cells exhibits an unusually shallow pH-dependence and that human LAMP1 expression enhances fusion at mildly acidic pH (Fig. 3B). This pH-dependence contrasts with the steep pH-dependencies of other viral fusion reactions (e.g., (44, 75)). Perhaps, acid-induced conformational changes in GPC occur in a less cooperative manner in the absence of LAMP1. Consistent with the ability of low pH to induce refolding of LASV GPC, exposure to pH 4.0 in the absence of target cells leads to conformational changes and GP1 shedding from GP2 (25, 33), resulting in functional inactivation of this protein (Suppl. Fig. S1 and (33)). This feature of LASV fusion is in contrast to Avian Sarcoma and Leukosis Virus Env glycoprotein, which becomes responsive to low pH and can mediate membrane fusion only after binding to a cognate receptor (44, 76). We note that, although our finding that LASV fusion can occur at mildly acidic pH is in general agreement with other reports (22, 31, 33), a few studies have reported a considerably lower pH range for GPC-mediated cell-cell fusion (34, 77). The reason for these divergent findings is unclear. We suggest that, because of the relative inability of GPC-mediated fusion pores to enlarge, the small fluorescent dye mixing-based fusion assay employed in our study more sensitively detects cell fusion than the syncytium- or reporter protein-based assays used by others.

We also characterized the nature of a fusion block imposed by ST-193, a pan-arenavirus fusion inhibitor targeting the interface between GP2 and the stable signal peptide of the GPC (62). At a ST-193 concentration that almost completely inhibits LASV GPC-mediated cell fusion, the compound still permits the fusion to proceed to a hemifusion stage. However, for the most part, these are not on-path hemifusion structures that naturally resolve into full fusion upon removing the inhibitor; their existence can only be revealed by CPZ treatment that destabilizes the hemifusion diaphragm. These results demonstrate that, under our experimental conditions, ST-193 does not block early pH-induced conformational changes in GPC. The tendency to form dead-end hemifusion structures may suggest a role for the stable signal peptide in late stages of fusion. Alternatively, non-productive fusion pathways can be favored by a less efficient GPC activation in the presence of ST-193.

Another striking feature of Old and New World arenaviruses, which use distinct cellular receptors for infection (1–3, 78), is that, in contrast to the overwhelming majority of enveloped viruses, arenavirus entry/fusion is resistant to restriction by IFITMs (35, 73, 74). We have recently shown that IFITM3 exerts broad antiviral activity by blocking fusion at a hemifusion stage through modulating the properties of the cytoplasmic leaflet of endosomal membranes (79). These findings may suggest that the GPC fusion machinery is more robust than other viral glycoproteins and is therefore able to overcome IFITM-mediated restriction. However, we have shown that: (1) as is the case for other viral proteins, GPC cannot overcome the increased membrane bending energy imposed by lyso-PC incorporation (Fig. 4B); and (2) LASV appears to avoid IFITM3 restriction by entering and fusing with endosomes devoid of this antiviral protein (73). It thus appears that New and Old World arenavirus trafficking pathways converge to unique cellular compartments devoid of IFITMs.

The key finding of this study is the discovery that LASV GPC-mediated fusion is enhanced by the late endosome-resident lipid, BMP. Since GPC-mediated fusion is not modulated by another anionic lipid, DOPS, the effect of BMP appears specific. Importantly, BMP modulates the late stages of LASV GPC-mediated cell fusion by promoting both the transition from hemifusion to full fusion and the growth of fusion pores (Figs. 6 and 7). BMP is an unusual lipid that is highly concentrated in late endosomes/lysosomes; it is involved in intracellular cholesterol transport and, through binding to Alix, in biogenesis of multivesicular bodies (65, 66, 70, 80, 81). Its intrinsic curvature is modulated by pH and calcium ions, as evidenced by the pH gradient-driven inward budding and low pH-dependent liposome fusion observed in BMP-containing liposomes (63, 80). Of note, the effect of exogenously added BMP on forced LASV pseudovirus fusion with plasma membranes is modest, and no detectable enhancement of viral entry through the endocytic pathway is detected. It is possible that exogenous addition of this lipid does not augment LASV pseudovirus fusion, probably because late endosomes contain optimal amounts of BMP.

The role of lipids in promoting viral fusion has been reported for alphaviruses (e.g., (82, 83)), flaviviruses (e.g., (84, 85)), orthomyxoviruses (86) and lentiviruses (48)). It should be stressed that, unlike the fusion of Dengue virus that is non-specifically promoted by anionic lipids (84), LASV fusion is specifically enhanced by BMP. We have also observed a similar effect of BMP on fusion of the Vesicular Stomatitis Virus G protein pseudotyped particles with supported lipid bilayers (87).

How does BMP specifically promote LASV fusion with late endosomes? Since this lipid appears to be present in both inner and cytoplasmic leaflets of endosomal membranes (68), it is feasible that BMP interacts directly with LASV GPC and augments functional oligomerization and/or refolding of this protein into the final 6-helix bundle structure. It is currently unclear whether BMP in the cytoplasmic leaflet may destabilize a hemifusion diaphragm and thereby promote the transition to full fusion. This possibility appears less likely, because one would expect to see a non-specific enhancement of fusion mediated by other viral glycoproteins, which is not observed in our experiments. The role of BMP in dilation of GPC-mediated fusion pores may explain its equally notable effect on fusion upon pretreatment of either effector or target cells (Suppl. Fig. S2B). It is expected that that this lipid incorporates into the fusion pore through either GPC-expressing or target cell membrane.

Together, our findings suggest the following model for LASV entry into cells. LASV fusion may be initiated in early LAMP1-containing endosomes, but here it can only culminate in the formation of a stable hemifusion intermediate. This intermediate resolves into full fusion after virus enters late endosomes enriched in BMP.

## Materials and Methods

### Cells and reagents

COS7 cells and HEK293T cells were maintained in Eagle’s Medium with glucose, L-glutamine, and sodium pyruvate, supplemented with 10% Cosmic Calf Serum (HyClone, Logan, Utah), penicillin/streptomycin (ThermoFisher, Waltham, MA Cat# 15140-122). The quail QT6 and chicken DF-1 cells were obtained from ATCC (Manassas, VA). QT6 cultured in F-12K medium with 2 mM L-glutamine, 10% tryptose phosphate broth and 5% calf serum. DF-1 cells were grown in DMEM with 4.5 g/l glucose, L-glutamine, sodium pyruvate and 10% fetal bovine serum. COS7 cells grown to ∼60% confluency on a 35 mm culture dish were transfected with 4 μg of the plasmid expressing the Lassa virus GPC (Josiah strain, a kind gift from F.-L. Cosset (34) using a standard calcium phosphate method. Where indicated, HEK293T and QT6 cells were transfected with the plasmids expressing the wild-type LAMP1 or the LAMP1 D384 mutant (a gift from Ron Riskin (Weizmann Institute, Israel) (32), using a calcium phosphate method. LAMP1-mRFP plasmid was from Addgene (Cat# 1817).

The HIV-1-based packaging vector pR9ΔEnv was from Dr. Chris Aiken (Vanderbilt University). The pMM310 vector expressing BlaM-Vpr, psPAX2 lentiviral packaging vector, pcRev vector, pMDG-VSVG plasmid expressing VSV-G, H1N1 WSN HA and NA expression vectors were described previously (35, 88). The pNL4-3.Luc.R-E-packaging vector was obtained through the NIH AIDS Reagent Program (from Dr. Nathaniel Landau (89)).

#### Lipids

DOPC (1,2-dioleoyl-sn-glycero-3-phosphocholine), DOPS (1,2-dioleoyl-sn-glycero-3-phospho-L-serine), BMP (bis(monooleoylglycero)phosphate (S,R Isomer)) and lyso-PC (lauroyl-lysoPC) were obtained from Avanti Polar Lipids (Alabaster, AL). Chlorpromazine was purchased from Sigma (Cat# C8138). A soluble fragment of human LAMP1 was obtained from Origene Protein (Cat# TP720784). The LASV fusion inhibitor ST-193 was from Medchem Express (NJ, Cat.# HY-101441). Annexin V labeled with AF647 was obtained from Invitrogen (ThermoFisher, Cat# A232004). Mouse monoclonal anti-BMP antibody 6C4, neuraminidase and TPCK-treated trypsin were purchased from Millipore Sigma (Burlington, MA, Cat# MABT837, N2876 and T4376). Mouse IgG1κ isotype control was purchased from Invitrogen (ThermoFisher, Cat# 14-4714-85). Fatty acid-free BSA was from Sigma (St. Louis, MO, Cat# A1933).

### Pseudovirus production and single-cycle infection assay

The single-cycle infection-competent pseudotyped viruses were generated by transfecting HEK293T/17 cells grown to ∼75% confluency in 10-cm dishes with JetPRIME transfection reagent (Polyplus-transfection, Illkirch-Graffenstaden, France). The cells and plasmid transfection mix were incubated for 14 h at 37°C, 5% CO_2_, followed by incubation with fresh medium for another ∼34 h. The viral supernatants were filtered through 0.45 μm polyethersulfone filters (PES, VWR), aliquoted and stored at −80°C. The p24 content of pseudovirus preparations was quantified by ELISA, as described previously (90). To generate luciferase-encoding pseudoviruses, HEK293T/17 cells were transfected with 6 μg pNL4-3.Luc.R-E-, 0.5 μg pcRev and 4 μg Lassa-GPC or 2.5 μg each of WSN HA- and NA-expressing plasmids, respectively. For the BlaM assay, HEK293T/17 cells were transfected with 4 μg pR9ΔEnv, 2 μg BlaM-Vpr, pcRev and Lassa-GPC or WSN HA/NA as above. To produce shRNA encoding pseudoviruses, HEK293T/17 cells were transfected with 1.5 μg pMDG-VSVG, 3 μg psPAX2, and 4 μg pooled shRNA plasmids.

For the infectivity assay, target cells (∼2×10^4^ cells/well in black-clear bottom 96-well plates) and virus (0.2 ng p24/well) were centrifuged at 1550×*g* for 30 min at 4°C. Unbound virus was washed off, 75 μL/well of growth medium was added, and cultured at 37°C, 5% CO_2_. Forty-eight hours post-infection, the samples were incubated for 5 min at room temperature with equivalent volumes of Bright-Glo^TM^ firefly luciferase substrate (Promega, Madison, WI), and the luciferase signal was measured using a TopCount NXT plate reader (PerkinElmer Life Sciences, Waltham, MA, USA).

### shRNA knock down and Western blotting

For shRNA knock down, 5 validated LAMP1 shRNAs were ordered from human Mission lentiviral library (Sigma). HEK293T/17 and A549 cells grown to ∼70% confluency in 6-well plates were transduced with 0.5 ng p24/well of shRNA pseudoviruses, with centrifugation at 1550×*g* for 30 min at 4°C. Unbound viruses were removed, fresh medium was added, and samples were incubated at 37°C, 5% CO_2_. Twenty-four hours post-transduction, cells were transferred to 10-cm dishes in the presence of 1.5 μg puromycin. After 4 days of selection, cells were processed for Western blotting, as described in (73). The LAMP1 protein was detected with rabbit anti-LAMP1 (Sigma, Cat# SAB3500285) and horseradish peroxidase-conjugated mouse anti-rabbit (Millipore, Cat# AP188P), using ECL Prime chemiluminescence reagent (GE Healthcare). The chemiluminescence signal was detected using a XR^+^ gel doc (Bio-Rad). Densitometry analysis was done using Image Lab software (Bio-Rad).

### Cell-cell fusion assay

Effector and target cells were labeled and cell-cell fusion quantified, as described in (46). Briefly, ∼2×10^6^ COS7 and target (HEK293T or avian cells, as indicated) cells were loaded with 1.5 μM calcein-AM or 20 μM CMAC (Invitrogen), respectively, for 30 min at 37°C. Labeled effector COS7 cells and target HEK293T or QT6 cells were detached from culture dishes using a non-enzymatic solution (divalent-free PBS/EGTA/EDTA). Effector and target cells were resuspended in PBS^++^ or PBS^++^ supplemented with 1 mg/ml BSA, as indicated, mixed at an 1:1 ratio in a test tube and added to wells of an 8-well slide (Thermo Fisher) coated with poly-lysine (Sigma, Cat# P1274). Cells were allowed to adhere and establish contacts for 30 min at room temperature. The density of cells was adjusted to minimize aggregation and, at the same time, ensure sufficient cell-cell contacts to allow subsequent fusion. Next, the pH was lowered for 10-20 min at room temperature or at 37°C to the indicated value, using a citric acid-sodium citrate buffer. Following the acid trigger step, cells were returned to pH 7.2 PBS^++^ and further incubated at 37°C for 30-60 min. Cell-cell fusion was detected by fluorescence microscopy based upon the appearance of calcein- and CMAC-positive cells, as described previously (46). The extent of fusion was determined by normalizing the number of double-positive cells per field to the total number of effector/target cell contacts. Typically, 10 randomly selected image fields, each containing ∼10 effector/target cell pairs, were analyzed for each well.

### Arresting fusion intermediates and testing the effects of exogenous lipids

To reveal the presence of unresolved hemifusion structures after triggering fusion of effector and target cells by exposure to low pH, cells were treated with 0.5 mM CPZ in PBS^++^ for 1 min at room temperature. To capture cells at a cold-arrested stage (CAS), effector/target cell pairs were chilled on ice and exposed to pH 5.0 at 4°C for 10 min. Cells were returned to PBS^++^ and either incubated at 37°C to allow fusion or treated with 0.5 mM CPZ for 1 min and immediately counted under the microscope.

To capture cell-cell fusion at a lipid-arrested stage (LAS), effector/target cell pairs were pretreated with 285μM stearoyl lyso-PC in PBS for 30 min at room temperature followed by exposure to a pH 5.0 buffer for 10 min at room temperature in the presence of lyso-PC. Cells were returned to neutral pH and either incubated at 37°C in the presence of lyso-PC or washed with delipidated BSA (2 mg/ml, Sigma) to remove lyso-PC and then maintained at 37°C for 30 min.

To assess the effect of lipids on cell fusion, COS7 cells mixed with target QT6 or HEK293T cells were allowed to adhere to a poly-lysine coated slide for 30 min at room temperature. Lipids DOPC, DOPS or BMP were diluted from 5 mg/ml stock solutions in ethanol to a final concentration of 10 or 40 μg/ml in PBS^++^ supplemented with 1 mg/ml BSA (Sigma) and immediately added to cells. Cells were incubated with lipids for 20 min at room temperature and exposed to acidic buffers at the pH and temperature indicated. Cell-cell fusion was scored after returning the cells to neutral pH and incubating for additional 30 min at 37°C. The effects of exogenous lipids on the late steps of LASV fusion were assessed by capturing the fusion reaction at CAS (exposure to a cold pH 5.0 buffer for 10 min) and adding 10 μg/ml of indicated lipids at neutral pH and incubating for 10 min at room temperature. Cells were then shifted to 37°C and incubated 30 min to allow fusion to occur.

### Pore permeability measurements

Permeability of pores between effector and target cells was measured, as previously described (71, 72). Briefly, GPC-transfected COS7 cells were loaded with calcein-AM dye, premixed with target HEK293T cells, and allowed to adhere to poly-lysine coated coverslips for 30 min at room temperature. Pieces of coverslip with cells were transferred into a home-made imaging chamber with an IR-absorbing coverslip at the bottom; the chamber contained a pH 5.0 buffer.

Temperature was quickly and locally raised and maintained at 37°C by illuminating the absorbing coverslip with an IR diode (71, 72). Dye transfer was monitored with a Fluoview300 laser scanning confocal microscope (Olympus IX70, America, Melville, NY) using an UPlanApo 60X/1.20NA water-immersion objective. Total fluorescence intensities of effector and target cells over time were determined using two manually drawn ROIs, and the relative pore permeability was calculated using the equation: *P*(*t*) = *Q*(*t*)·(*dI_t_*/*dt*)/(*I_e_*(*t*) - *I_t_*(*t*)) = −*Q*(*t*)·(dI_e_/dt)(*I_e_*(*t*) - *I_t_*(*t*)), where I_t_ and I_e_ are fluorescence intensities of target and effector cell, respectively, and Q(t) is a correction factor to account for the temperature-dependence of the diffusion coefficient of calcein (71). To assess the effect of exogenous lipids, pieces of a coverslip with effector and target cells were pretreated with BMP or DOPC (20 μg/ml), as described above, for 10 min at room temperature, prior to transferring the coverslips to the imaging chamber containing a pH 5.0 buffer; temperature-jump experiments were then performed.

### Virus-cell fusion assay

Target cells were seeded in 96-well plates one day before the experiment to reach ∼90% confluency the next morning. Cells were pretreated with 0.2 μM Bafilomycin A1 (Santa Cruz Biotech., Cat# SC201550) for 1 h at 37°C. Cells were then spinoculated with various amounts of LASV, JUNV or IAV pseudoviruses containing BlaM-Vpr in HBSS supplemented with 10% FBS at 3,000 rpm for 30 min at room temperature. Cells were washed to remove unbound viruses, treated with 40 μg/ml of indicated lipids dissolved in PBS++ supplemented with 1 mg/ml BSA for 10 min at room temperature and exposed to a pH 5.0 buffer for 30 min at 37°C. Cells were loaded with the BlaM CCF-4-AM substrate (Thermo Fisher Scientific) and incubated overnight at 16°C. For the regular virus-cell fusion assay, cells were pretreated with lipids as described above, spinoculated with pseudoviruses and placed into a CO_2_ incubator for 2 h at 37°C. The resulting cytosolic BlaM activity was measured on a Wallac 1420 Victor2 (PerkinElmer, Turku, Finland) fluorescence microplate reader at 460nm (blue) and 528nm (green), using 400nm excitation.

For virus-cell fusion experiments involving antibodies, the target cells grown to ∼60% confluency in black-clear bottom 96-well plates were starved for 6 h and incubated for 15 h with 50 μg/mL antibodies in growth medium. Pseudoviruses containing BlaM-Vpr were bound to cells, as described above, and samples were incubated for 2 h at 37°C, 5% CO_2_. Cells were then loaded with CCF-4-AM substrate, as described above. The BlaM activity was measured using a SpectraMax i3X fluorescence plate reader (Molecular Devices, Sunnyvale, CA).

### Immunofluorescence/Cell surface staining

COS7 cells were treated for 20 min at room temperature with the indicated lipids that were dissolved at 10 μg/ml in PBS supplemented with 1mg/ml BSA. Cells then washed and fixed with 4% PFA (Electron Microscopy Sciences, Cat# 15710). The presence of DOPS on the cell surface was detected using Annexin V labeled with AlexaFluor647 (1:200 dilution of the stock). Exogenous BMP on the cell surface was detected using the mouse anti-BMP 6C4 monoclonal antibody followed by incubation with goat anti-mouse IgG conjugated with AlexaFluor488 (Invitrogen).

## Acknowledgements

The authors wish to thank F.-L. Cosset (University Claude Bernard Lyon) for the gift of the LASV GPC plasmid and Jack Nunberg (University of Montana) for the plasmid encoding wild-type and mutant GPC of Junin virus. We thank Gokul Raghunath and other members of the Melikyan laboratory for critical reading of the manuscript and constructive suggestions.

## Author Contributions

GBM and FSC conceived the study; RMM and MM performed the experiments; YZ provided reagents; RMM, MM and GBM analyzed the results; GBM obtained funding and supervised the project; GBM wrote the initial draft of the manuscript; all authors edited the manuscript.

## Supplemental Figure Legends

**Figure S1. LASV GPC inhibition by ST-193 and inactivation at low pH.** (A) Dose-dependent inhibition of LASV GPC-mediated cell-cell fusion by the arenavirus fusion inhibitor ST-193. LASV GPC-expressing COS7 cells (loaded with calcein-AM) and HEK293T cells transfected with LAMP1mut (loaded with CMAC) were mixed at a 1:1 ratio, adhered to poly-lysine coated coverslips and incubated for 30 min at room temperature. Cell fusion was triggered by exposure to pH 6.2 for 10 min at room temperature (suboptimal trigger) in the presence or absence of the indicated concentration of ST-193. The results are means and SEM from three independent experiments. (B) GPC-expressing COS7 cells were pretreated with a pH 4.0 buffer for 10 min at 37°C followed by co-incubation with target HEK293T cells for 10 min at neutral pH, room temperature, to establish cell-cell contacts. Effector-target cell complexes were then exposed to pH 6.2 for 10 min at room temperature and further incubated at neutral pH for 1 h at 37°C. The results are means and SEM from three independent experiments.

**Figure S2. Incorporation of exogenously added lipids into COS7 cells.** (A) *Top*: Representative images of Annexin V-stained cells. Cells were pretreated with 10 μg/ml DOPC (negative control) or DOPS in BSA for 20 min at room temperature, washed and stained with 5 μg/ml of Alexa-Fluor 647-labeled Annexin V (ThermoFisher) for 1 h at 4°C. *Bottom*: Cells were pretreated with 10 μg/ml BMP in BSA or mock treated, washed, fixed with 4% PFA and incubated with anti-BMP antibody(1:250) for 1h at 4°C, followed by incubation with goat anti mouse Alexa-Fluor 488-conjugated second antibody (1:1000 dilution). Images were acquired on a Fluoview300 microscope (Olympus, Melville, NY), using an UPlanApo 60X/1.20NA water-immersion objective and standard eGFP and Cy5 filter cubes for Annexin V and BMP immunostaining, respectively. (B) Effect of BMP on cell fusion upon incorporation into target or effector cells. Either GPC-expressing COS7 cells or HEK293T cells were pretreated in suspension with either DOPC or BMP at 10 μg/ml for 20 min at room temperature. Cells were washed mixed with target or effector cells, respectively, and allowed to adhere to glass slides for 15 min at room temperature. Fusion was triggered by exposure to pH 5.0 for 20 min, 37°C. The results are plotted as fold-increase in cell-cell fusion by BMP relative to DOPC. Data are means and SEM from four independent experiments.

**Figure S3. Effect of exogenous lipids on LASV pseudovirus fusion.** (A, B) Two independent experiments showing the effect of exogenously added DOPC, DOPS and BMP (40 μg/ml in BSA-containing PBS) on LASVpp fusion with HEK293T cells. The LASVpp fusion under these conditions was measured by a BlaM assay using the blue/green ratio of fluorescence of a BlaM substrate that was loaded into cells. (C, D) Same as in panel A, but showing the result of two experiments of LASVpp fusion with avian DF-1 cells. No significant changes in the BlaM signal were observed after cell pretreatment with any of the lipids.

